# MP-GNN: Graph Neural Networks to Identify Moonlighting Proteins

**DOI:** 10.1101/2023.11.13.566879

**Authors:** Hongliang Zhou, Rik Sarkar

## Abstract

Moonlighting proteins are those proteins that perform more than one distinct function in the body. They are pivotal in various metabolic pathways and disease mechanisms. Identifying moonlighting proteins remains a challenge in Computational Biology. In this work, we propose the first graph neural network based models to identify moonlighting proteins. Our models work on large protein-protein interaction (PPI) networks with sparse labels of moonlighting and non-moonlighting proteins. In addition to PPI network, the models make use of features derived from the amino acid sequences of the proteins.

We propose two frameworks: one as graph classification based on the local neighborhood of the query protein; and the other node classification based on the entire graph. These GNN-based methods outperform traditional machine learning methods that have previously been used for moonlighting prediction. The global full network-based model, operating on *Homo sapiens* data achieves accuracy of 88.4% and F1 score of 88.8%. The local neighborhood method is more lightweight and can be applied to larger protein sets with multiple species.

**CCS CONCEPTS:** **• Applied computing → Computational proteomics.**

## 1 INTRODUCTION

In the 1980s, it was discovered that certain crystallin proteins occurring in eye lenses have amino acid sequences that match enzymes expressed in other tissues [29, 30]. Experiments confirmed that they act like the enzymes in metabolic reactions, thus demonstrating that the same proteins expressed from the same genes could perform very different functions depending on where it is expressed [26]. These dual function proteins were later given the name *Moonlighting proteins*. Since then, the canonical functions of moonlighting proteins have been often found to be associated with critical biological pathways such as Krebs cycle or glycolysis. They are also frequently associated with pathogen virulence [3]. Thus, identification and understanding of moonlighting proteins is important for fundamental understanding of biology, of bacterial activity and drug design [10].

Moonlighting proteins (abbreviated MPs) have been discovered in all kinds of living organisms, but the experimental methodologies to find proteins with multiple functions remain challenging and expensive. According to MoonProt [20] – a database for Moonlighting proteins – only about 500 MPs have been identified to date. Thus, even with our knowledge of billions of protein sequences found by current high throughput technologies [22], we need fundamental developments in understanding MPs and in methods for finding them. Computational techniques that identify good candidates to test for moonlighting properties can narrow the field for experimentation and significantly reduce the cost and effort in finding new moonlighting proteins.

### Related works

Statistical machine learning methods have been developed for identification of moonlighting proteins. Methods such as MPFit [14] and DextMP [13] have achieved good accuracy by making use of Gene Ontology (GO) and natural language processing based on literature context respectively. However, these features require human analysis and labelling, making them unsuitable for all but a few select proteins. For example, only a small fraction (≈ 0.1%) of proteins in the Uniprot database carry GO annotations. More recent approaches have made use of more universal features – amino acid sequences and Protein-Protein Interaction (PPI) networks. Features derived from amino acid sequences have formed the basis of conventional machine learning models including Deep neural networks (DNNs), Support Vector machine (SVM) and Random Forest (RF) [25]. K-nearest neighbors was investigated in Hu and Du [9] context of MPs. By definition, these models base their predictions on protein structure and ignore interactions. In contrast, PPI based methods base their predictions purely on inter-protein interactions [12, 14]. These methods used statistical measures and bioinformatic techniques, including remote homology searches, co-evolution, and network analysis, to explore the interactomic features from PPI interactions [11, 7, 8, 12, 14]. However, this approach requires a significant amount of domain knowledge and human expertise. Moreover, these networks are not directly compatible with conventional machine learning models.

### Our contributions

Graph neural networks have been developed as a class of machine learning models that can learn from graphs as well as vector features (See [36] for a survey). They have been applied to a range of computational biology problems including protein function prediction, structure prediction and PPI link predictions [11, 17, 31–33, 35].

In this paper, we present the first graph neural network (GNN) framework MP-GNN – designed to identify moonlighting proteins. The models are trained on a PPI network, where the nodes are labeled with sequence-based features. We study the impact of important graph neural network concepts: Graph convolutions [16], Graph attentions [28] and GraphSAGE sampling [6] are compared. The neural networks are designed for two different cases (Section 3). First, we introduce Global MP-GNN. When the PPI network consists of proteins of a single species, we devise this global version of MP-GNN which makes use of the full PPI network interconnectivity information, and approach the MP prediction as a node classification task. When analyzing larger networks such as ones with proteins from all species, this approach is impractical and we devise a local version of MP-GNN which is trained on individual node neighborhoods and approach the MP prediction as a graph classification task. To overcome the challenge of small number of labelled proteins, we incorporate a semi supervised learning technique that is found to improve accuracy.

Experiments (Section 4) on the *Homo sapiens* data demonstrate that global MP-GNN outperforms previous methods in terms of accuracy, F-1 score, and precision. The Local MP-GNN models surpass previous methods. The GraphSAGE model with a self-attention pooling (SAGPooling) layer achieved the best performance, with an accuracy of 79.5%, an F-1 score of 79.0%, a precision of 80.0%, a recall of 78.0%, and a ROC-AUC of 75.8%. Using semi-supervised learning, the model achieved even better performance, with an accuracy of 88.4%, a recall of 87.2%, an F-1 score of 88%, a precision of 88.8%, and a ROC-AUC of 81.3%.

Investigating the edge weights predicted by the Graph Attention Network (GAT) model on local ego-networks showed that most self-loops are assigned larger weights, suggesting that GAT prefers to preserve the distribution of weights in neighborhood.

In the following, Section 2 summarizes the basics of graph neural networks and its variants. Readers familiar with these topics may wish to skim it and move ahead to the next section.

## 2 TECHNICAL BACKGROUND: GRAPH NEURAL NETWORKS

Graph neural networks work by simulating message passing and aggreation between nodes. Suppose 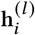 is the feature vector of node *i* at layer *l*, then the core of GNNs is to implement the following type of function:

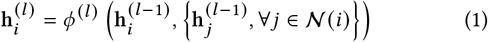

where 𝒩 (*i*) is the set of neighbours of the node *i, ϕ* is some aggregate and combine function. That is, at every message passing layer, the features of a node is updated to an aggregate of the features of the previous layers. Through one layer, the features at a node becomes an aggregate of its neighborhood. Over multiple message passing layers, it becomes an aggregate of a multi-hop neighborhood. Depending on the application, the GNN also contains fully connected layers, poooling layers etc. to achieve specific objectives like classification.

### 2.1 Graph Convolutional Network (GCN)

Graph convolutional networks [16] are a variant of GNNs with a class of layer functions given by:

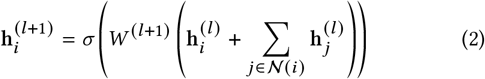

where *W* is a learnable weight matrix, and *σ* is a non-linear activation function such as ReLU. The notable feature here is that the aggregation is linear over the nighborhood, but the fetaure weights are modified by *W*, followed by the non-linear activation *σ*.

### 2.2 Graph Attention Network (GAT)

Unlike GCNs, GATs assign different importances to different edges (or neighbors) [28]. The layers perform the following function:

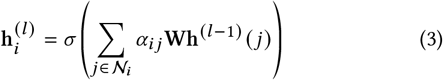

where *α*_*i j*_ is a softmax function:

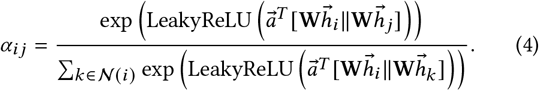

Here 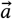 is a vector of learnable weights and ∥ represents vector concatenation. The coefficient *α*_*i j*_ represents the importance of node *i* to node *j*.

### 2.3 GraphSAGE (SAmple and aggreGatE)

In many graph problems, the number of edges can be large resulting in large sizes for neighborhoods 𝒩_*i*_. Methods such as GCN are also mainly transductive, meaning they do not perform well on unseen data. E.g. they do not cope well with graphs or nodes not seen in training. GraphSAGE [6] introduced sampling to reduce data complexity an improve generality. The update function of GraphSAGE is the similar to GCN, except that the neighbourhood is a smaller set randomly drawn from the real neighbours 𝒩_*i*_. GraphSAGE also performs better on unseen nodes.

## 3 MP-GNN: NEURAL NETWORKS FOR MOONLIGHTING PROTEIN PREDICTIONS

In this section, we first describe the label features we use to enhance model performance, then the construction of the PPI networks and finally the design of neural networks and their training process.

### 3.1 Sequence derived physiochemical features

Physical and chemical properties of proteins are determined by their amino acid sequences. Thus features derived from these sequences are called structure derived physiochemical features, or *physiochemical features* in brief. Formally, a physiochemical feature can be seen as a function *f* : 𝒫 → ℝ^*n*^ – embedding any sequence in the set of possible sequences 𝒫 into a Euclidean space.

Such physiochemical features have been the basis of using machine learning methods like KNN, RF, and SVM [2, 9, 25]. In this paper we make use of use the following physiochemical features that were introduced in [2].

#### 3.1.1 quasi-sequence-order (QSOrder)

The quasi-sequence-order (QSOrder) feature, denoted as *f* (*k*), is defined based on the normalized occurrence of amino acid type *k* (where *k* ranges from 1 to 50) The formula for *f* (*k*) is given by:

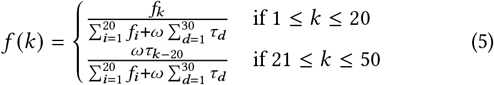

In this formula, *f*_*k*_ is the frequency of amino acid type *k. ω* is a weighting factor and is given as 0.1 in our setting, and τ_*k*_ is the Sequence-order coupling number, defined in [2]. This descriptor provides a combination of sequence composition and sequence order information.

#### 3.1.2 Amino Acid Composition (AAC)

Amino Acid Composition (AAC) refers to the proportion of each amino acid in a protein sequence, for the 20 sommon amino acids. Specifically, the AAC for a protein sequence 𝓅 is a vector in ℝ^20^, in which the *i*^*th*^ entry of the AAC vector *x* is calculated as:

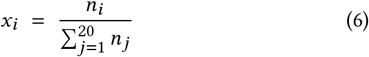

where *n*_*i*_ represents the number of occurrences of amino acid *i*.

### 3.2 Full PPI networks and Global MP-GNN model

#### 3.2.1 Construction of Full PPI networks

Each protein in the graph *G* = (*P, A*) is represented as a node included in the set *P*. For each pair of proteins *P*_*i*_ and *P* _*j*_, a confidence value *e*_*i,j*_ represents the likelihood of interaction (known from existing experimental or statistical data). We add an edge A_*i,j*_ = 1 if the confidence *e*_*i,j*_ *> θ*, where *θ* is a predetermined threshold. When they do not interact or the edge comes with a confidence lower than the threshold, we set A_*i,j*_ = 0.

#### 3.2.2 Global MP-GNN design and pipeline

Figure 1(a) illustrates the pipeline of the global MP-GNN framework. The GNN model is a trained model with a width of 256 and a depth of 3.

**Figure 1:**
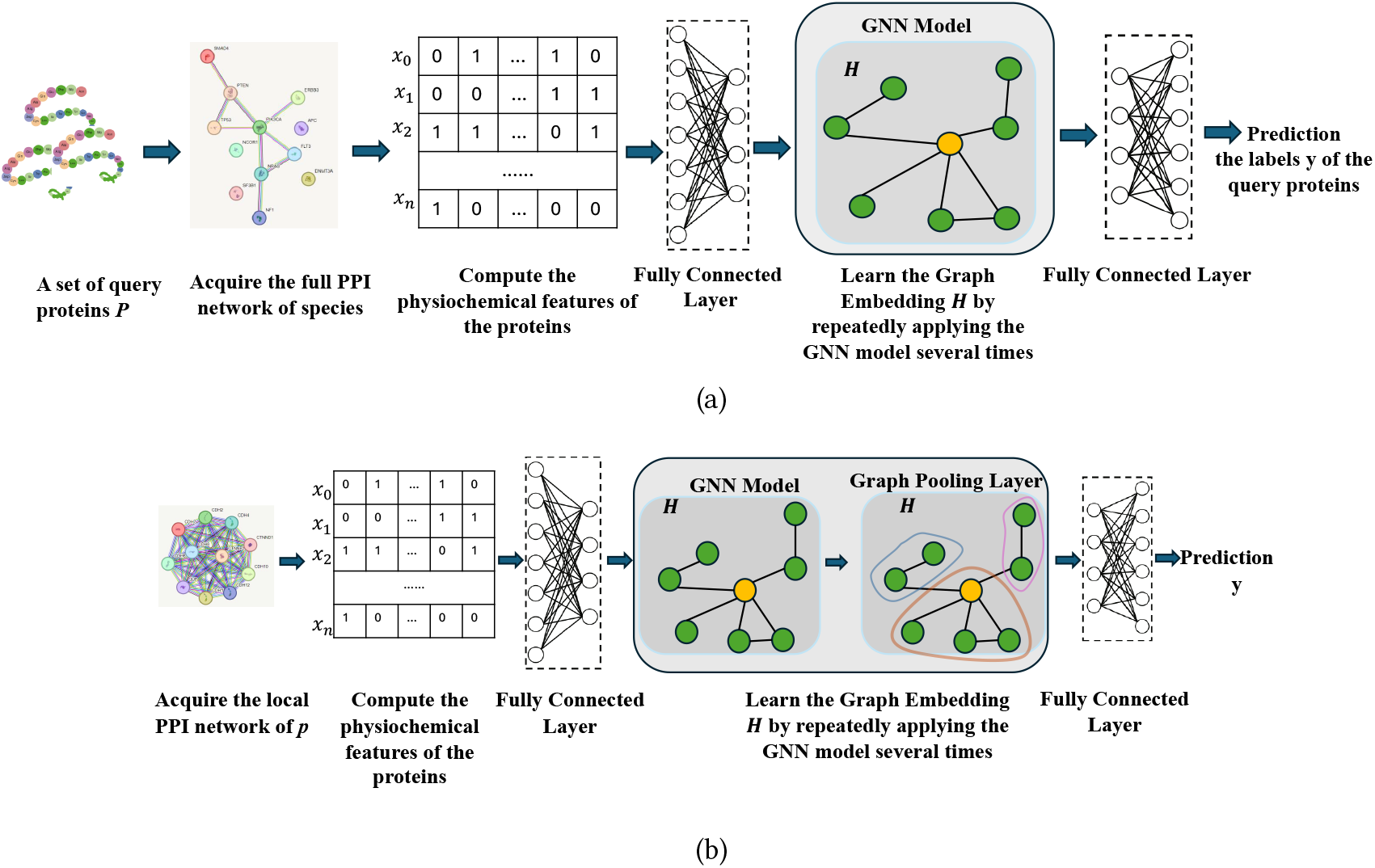
**(a) Global MP-GNN framework. Operating on the full PPI network of a species (e.g. Homo sapiens). The GNN has three layers and a width of 256. The GNN layers are repeated 3 times. (b) Local MP-GNN framework. Operating on the local neighborhood of a node. The GNN and pooling layers are repeated 3 times.**

Given a set of query proteins, we pass them through the following process: (i) Construct the partially labeled full PPI network *G* = (*P*, A, *y*) for any species as described above, each protein 𝓅 ∈ *P* can be classified as either an MP, with a label *y*_𝓅_ = 1, or a non-MP, with a label *y*_𝓅_ = 0. Only a subset of nodes, *P*_*L*_ ⊆ *P*, are labeled, and the unlabeled nodes are denoted by *P*_*U*_; (ii) extract the physiochemical feature *x*_𝓅_ of each protein 𝓅 ∈ *P*, and construct the node feature matrix X; (iii) apply the GNN model to predict label *y*_𝓅_ for each protein 𝓅 based on X and A.

### 3.3 Ego Networks and Local MP-GNN model

#### 3.3.1 Construction of local PPI networks

This approach uses the local neighborhood of a query protein to make the prediction. Given each protein we use its *K* closest interacting partners, to construct the local neighborhood graph called the ego network. We recommend setting the graph size between 20 and 100 closest interacting partners to balance the representation of interacting patterns against the inclusion of noise and uncertain edges. We assume all interactions are bidirectional. Like the full-network method, edges can be labeled with weights, and we apply a threshold *θ* to filter edges. For all pairs *i, j*, we set A_*i,j*_ = 1 if the confidence is greater than *θ*, and 0 if unconnected or below *θ*. Given each protein we use its *K* closest interacting partners, to construct the local neighborhood graph called the ego network. We recommend setting the graph size between 20 and 100 closest interacting partners to balance the representation of interacting patterns against the inclusion of noise and uncertain edges. We assume all interactions are bidirectional. Like the full-network method, edges can be labeled with weights, and we apply a threshold *θ* to filter edges. For all pairs *i, j*, we set A_*i,j*_ = 1 if the confidence is greater than *θ*, and 0 if unconnected or below *θ*.

#### 3.3.2 Local MP-GNN design and pipeline

An illustration of the pipeline is presented in Figure 1(b). We pass the graph into the GNN layer first, with a width of 256, and then to the graph pooling layer. We repeat this process three times. Graph pooling involves a form of coarsesning of the graph, where groups of nodes are aggregated into single nodes resulting in a more compact representation of the graph. There are several forms of graph pooling that can be used to coarsen graphs [36].

Given a query protein *p*, we pass it through the following process: (i) extract its *ego-network* with its closest *K* neighbours from the STRING database, as described in Section 4.1.3; (ii) construct the egonetwork **G**_**p**_ = (*V*_𝓅_, **A**_**p**_) of *p* as described above, extract the physio-chemical feature *x*_*p*_^′^ of each protein *p*^′^ ∈ *V*_*p*_, and construct the node feature matrix X; (iv) apply GNN model *GN N* : (**X**_**p**_, **A**_**p**_) → {0, 1} to predict whether the query protein is an MP.

### 3.4 Training

For each graph convolutional layer—GAT, GCN, and GraphSAGE— we adopted a three-layer structure with a width of 256. We employed *EarlyStopping* as regularization setting a patience of 30. The learning rate was dynamically adjusted, starting at an initial rate of 0.001 and halved after 10 epochs without performance gains. We used the Adam [15] optimizer, and the Binary Cross Entropy (BCE) loss [23] as the loss function:

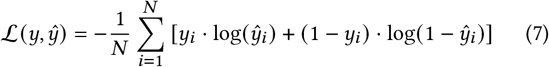

For model selection, we first randomized the data to prevent model bias. Then, we set aside 15% of the labeled data as an independent test set. The remaining labeled data underwent 5-fold cross-validation. In each fold, we trained the model for a maximum of 100 epochs on the training set, while monitoring the model’s performance with EarlyStopper on the validation set. Finally, we selected the top three models based on their performance on the validation dataset to form a soft-voting ensemble model. We then tested the performance of the ensemble model on the test set and used its performance to evaluate the model performance.

Then, to further enhance the model performance and address the challenge of sparse labeling of MPs and non-MPs, we applied *semi-supervised learning* [18]. Specifically, we utilize the *pseudolabel* method[21]. We first trained the models on the labeled data for 50 epochs, then we employed the models to predict unlabeled nodes and selected high-confidence predictions, with a threshold of 0.9, as *pseudo-samples*. Then, we augmented the labeled dataset with this *pseudo-samples* and continued training for an additional 30 epochs. This process was repeated for 10 times, and we recorded the final model performance on the test sets.

## 4 EXPERIMENTS

We carried out experiments in classifying moonlighting and non moonlighting proteins. We compare the various graph neural network ideas with conventional machine learning methods of KNN, SVM, RF and DNN found in [9, 25].

### 4.1 Dataset and network construction

This section describes the protein data used in experiments and the construction of PPI networks.

#### 4.1.1 Moonlighting Proteins

The experiments employed the same benchmark dataset as [25]. It consists of 351 proteins, including 215 moonlighting proteins (MPs) and 136 non-moonlighting proteins (non-MPs)^1^. These proteins represent a wide array of organisms, including humans, E. coli, yeast, rats, fruit flies, and thale cress. The statistics of the data are given in Table 1. The MP data were sourced from the MoonProt database [20]. To minimize redundancy and ensure data quality, proteins with more than 40% sequence similarity using CD-hit [4] are excluded. The non-MP examples were the same as those used in MPFit [14].

**Table 1:**
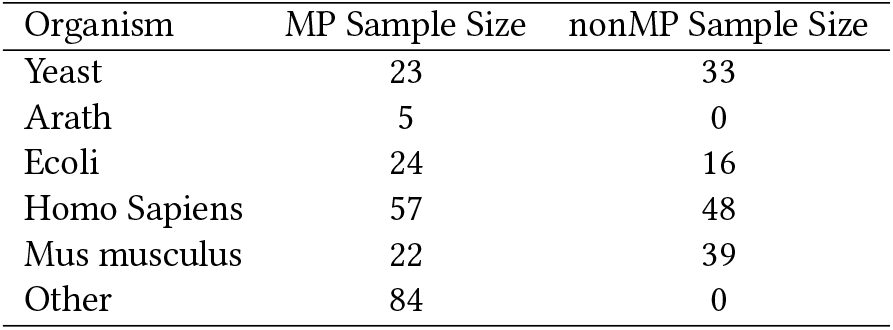
**Statistics for the data. The first column indicates the organism name, the last two columns indicate the contained MPs and non-MPs of the corresponding species in Shirafkan et al. [25]’s dataset.**

| Organism | MP Sample Size | nonMP Sample Size |
| --- | --- | --- |
| Yeast | 23 | 33 |
| Arath | 5 | 0 |
| Ecoli | 24 | 16 |
| Homo Sapiens | 57 | 48 |
| Mus musculus | 22 | 39 |
| Other | 84 | 0 |

#### 4.1.2 Global PPI Networks

To construct the global PPI networks, we acquired the full networks specifically for *homo sapiens*, since only the *homo sapiens’* MP and non-MP data are balanced and large enough(∼ 50). As of the latest update, the network encompasses 19,622 nodes, with interactions assigned a confidence value ranging from 0 to 1000. To balance data quality with graph size, we applied a confidence threshold of 250, resulting in a refined graph that contains 13,524,114 edges. Within this network, all 57 MPs and 48 non-MPs of *homo sapiens* were accurately matched.

#### 4.1.3 Local PPI Networks

We downloaded the Protein-protein Interaction (PPI) information from the STRING database, which provides comprehensive protein interaction data for each species [27].

To construct the local PPI networks, we queried the closest 10-100 interacting partners for each MP and non-MP. These interactions could be either physical or functional, and are assigned a score between 0 to 1 based on various types of evidence. To maintain interaction, we filtered these data by a threshold of 0.7. Out of the 215 MPs listed in Shirafkan et al. [25], 174 were found within the STRING database, accounting for 80.9% of the MPs. All example non-MPs were also successfully located within the database. Consequently, our finalized dataset comprised 310 proteins, including 174 MPs and 136 non-MPs. For each protein, we obtained PPI networks of various sizes, ranging from their 20 closest neighbors up to 100, to facilitate a thorough analysis.

Table 2 and Figure 2 present the results of various GNN models, in local and global, the supervised and unsupervised settings, and a comparison with four top-performing baseline methods as identified in the works of Shirafkan et al. [25] and Hu and Du [9]: KNN, SVM, RF, and DNN, on the data of *Homo Sapiens* only. As seen in Figure 2, while some conventional methods do well in some metrics only the graph neural networks do well consistently in all the metrics.

**Table 2:**
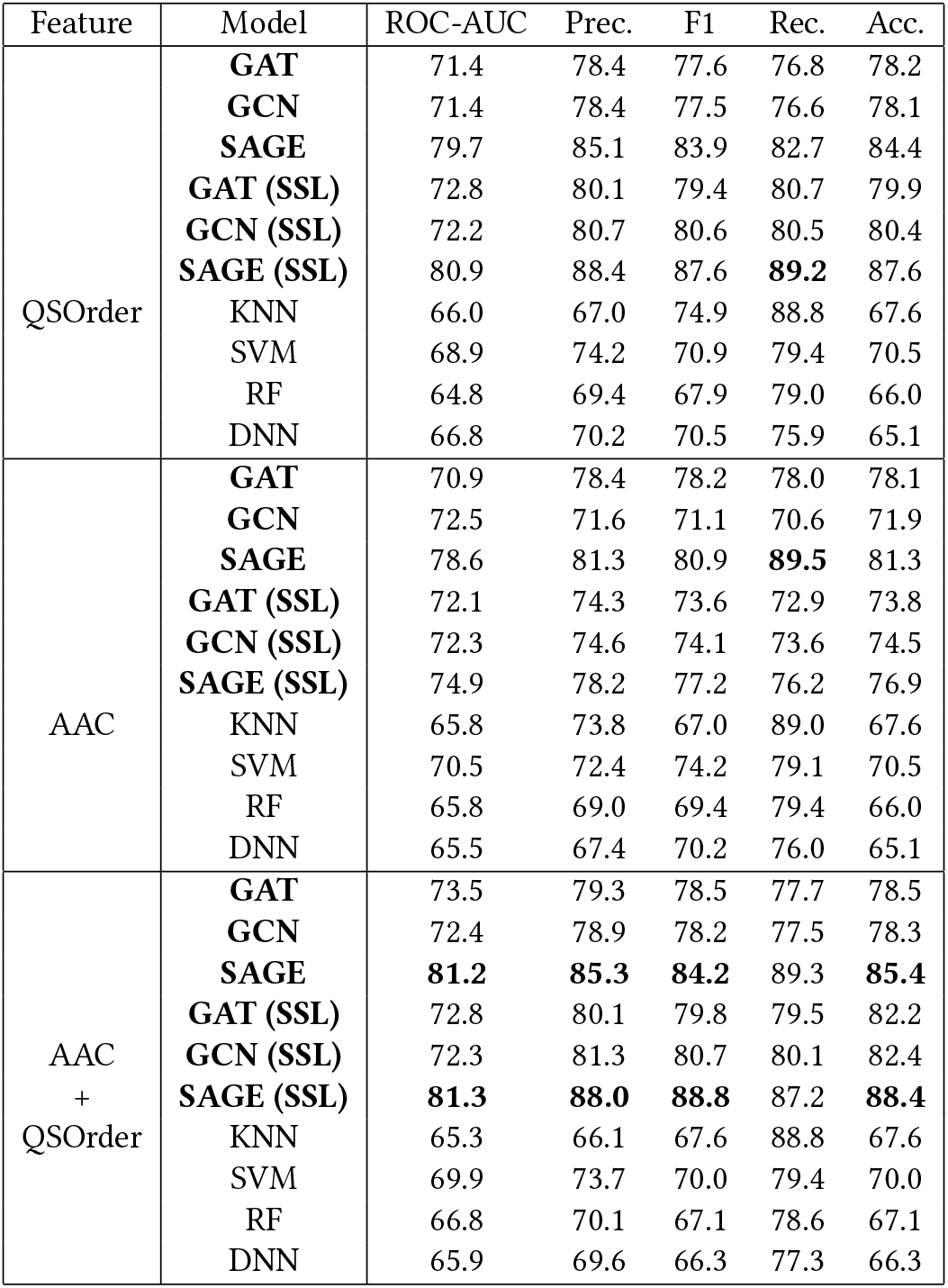
**The experimental results of different combinations of naive graph convolutional layers node features in the global MP-GNN framework, and the corresponding performances of the baseline models. The table shows that the global MPGNN models consistently outperformed previous methods; specifically, the graphSAGE model in the semisupervised learning setting achieved the top performance in all metrics, significantly outperforming previous methods by about 10-20%.**

**Figure 2:**
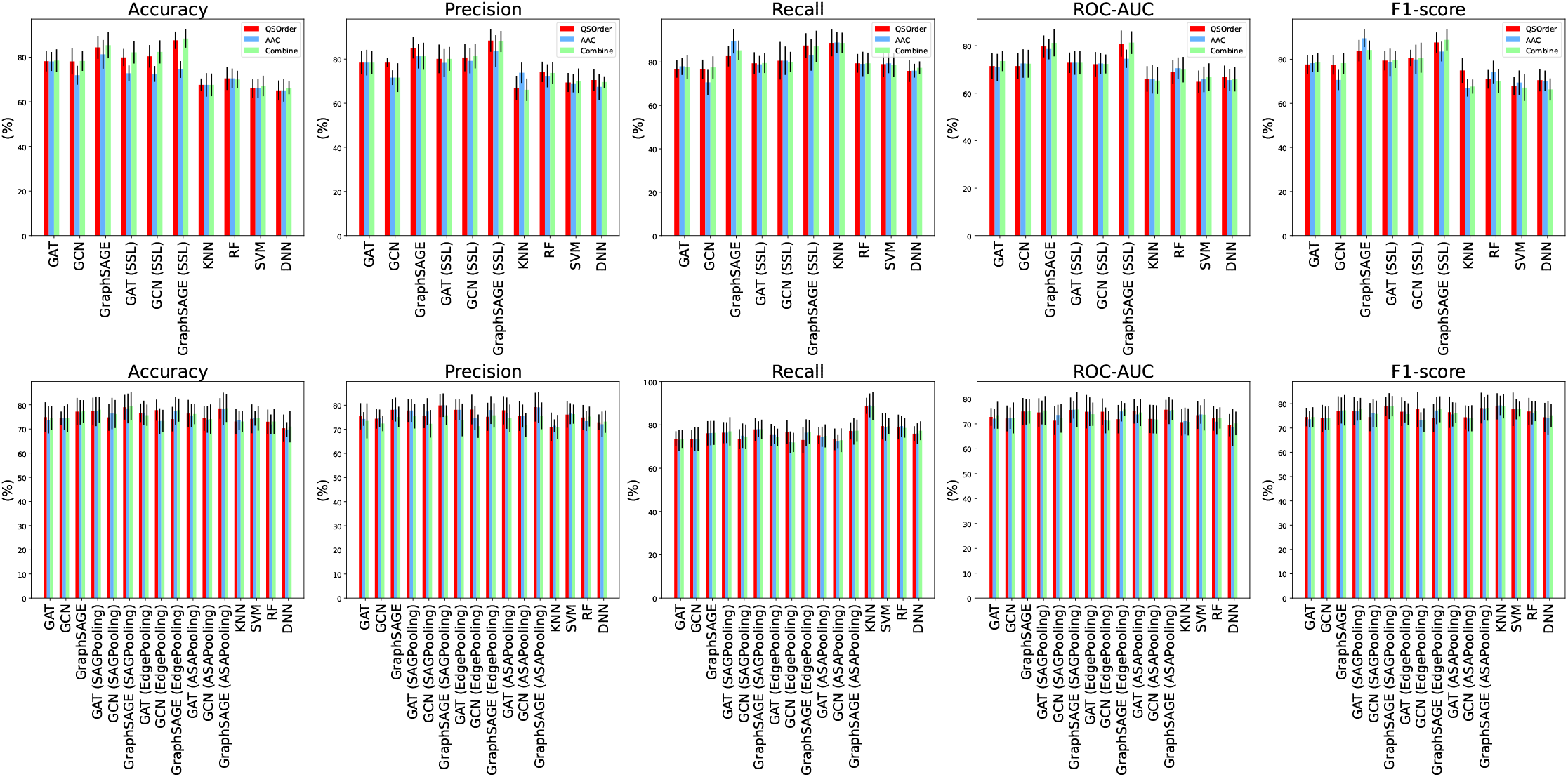
**A comparison between all global MP-GNN models (top row) and local MP-GNN models with different graph pooling layers (bottom row) and the baseline models. From the plots, we can see that the global MPGNN model with GraphSAGE in the semi-supervised learning setting exhibited the top performance, significantly outperforming other (both global and local) MPGNNs and the baseline models**

Generally, semi-supervised learning (SSL) is found to improve the model performance by about 2-3%, for all feature types. GraphSAGE exhibits the best performance across all metrics, in both the supervised and semi-supervised settings, and its performance is maximized with the AAC+QSOrder feature combination. In the supervised setting, the GraphSAGE model exceeded the top-performing baseline model by 11.1% in accuracy, 4.9% in F-1 score, 8.8% in precision, and 7.4% in ROC-AUC; in the semi-supervised setting, the model performance was further improved by about 2-3%, achieving an accuracy of 88.4%, recall of 87.2%, F1-score of 88.0%, precision of 88.8%, and ROC-AUC of 81.3%.

The results of GAT and GCN models were consistently below GraphSAGE by about 5% but still exhibit improvements of about 1-5% in all metrics over the baseline methods. Similarly, when we applied SSL to them, these models were improved by about 2-3% across every metric.

For features, we found that the combination of AAC and QSOrder produces the best results. Using GraphSAGE as an example, its accuracy with QSOrder alone was 84.4%, recall was 82.7%, F-1 score was 83.9%, precision was 85.1%, and ROC-AUC was 79.7%; its accuracy with AAC alone was 81.3%, recall was 89.5%, F-1 score was 80.9%, precision was 81.3%, and ROC-AUC was 78.6%. However, when using the concatenation of AAC and QSOrder as the node feature, its accuracy increases to 85.4%, F-1 score to 84.2%, precision to 85.3%, and ROC-AUC to 81.2%. Only recall did not show improvement. Similar observations were observed across all models, including previous methods.

### 4.2 Local MP-GNN

#### 4.2.1 Full Dataset Results

Table 3 presents the results for the local MP-GNN framework on various GNN variants on the full dataset, with various types of graph pooling layers. We can see that even the naive implementations of the GNN models have already slightly outperformed the baseline methods, and the combination of various graph pooling layers consistently further improved these models’ performance.

**Table 3:**
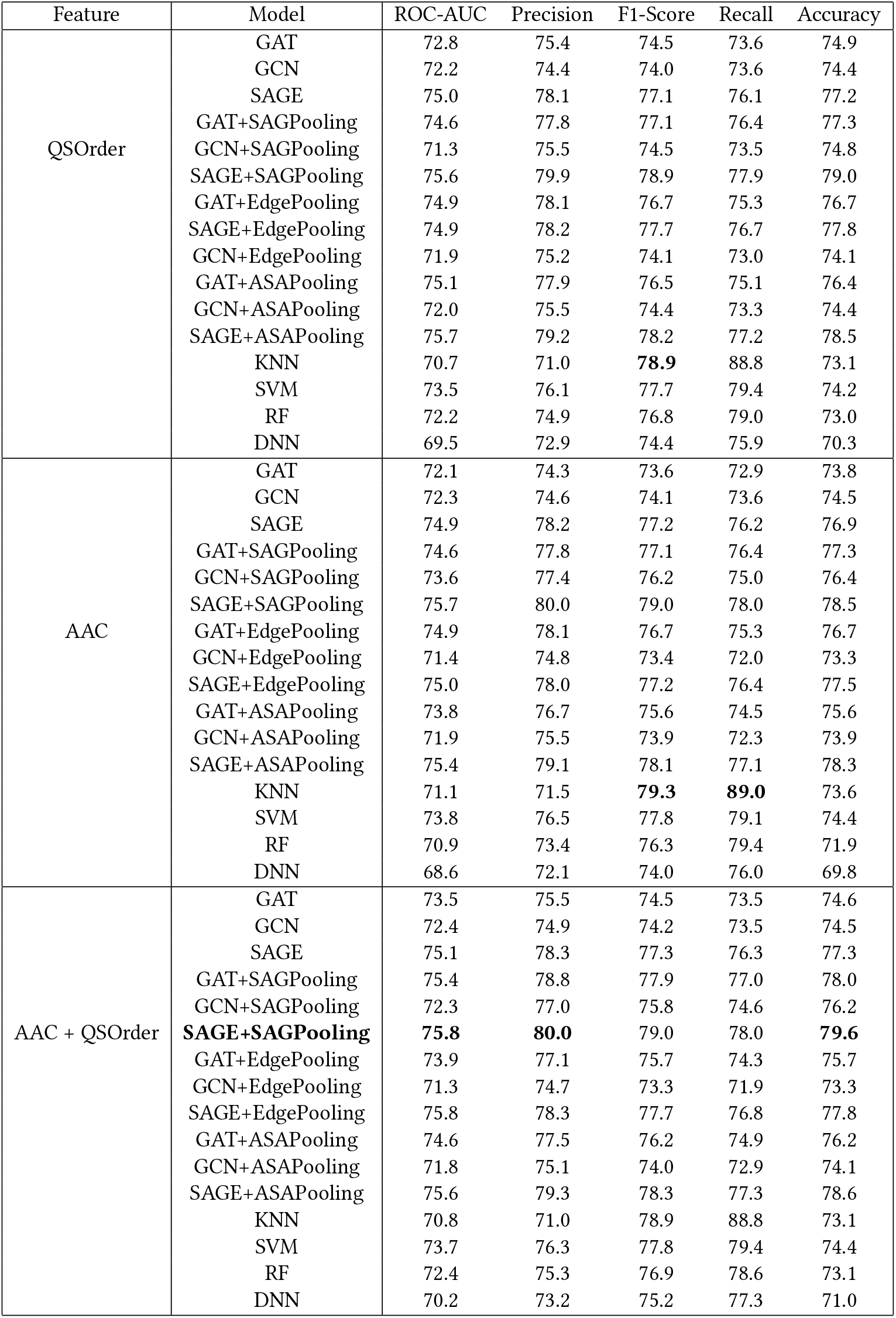
**The experimental results of different combinations of Graph Pooling Layers, naive graph convolutional layers node features in the local MP-GNN framework, and the corresponding performances of the baseline models. The table shows that graph pooling layers consistently improve the model performances. Specifically, the SAGPooling layer exhibited the best effect, and the combination of GraphSAGE, AAC+QSOrder, and SAGPooling demonstrated the best accuracy, ROC-AUC, and Precision.**

The GraphSAGE model with SAGPooling layer, utilizing the concatenation of QSOrder and AAC achieved the best performance, with accuracy of 79.6%, recall of 78.0%, F1-score of 79.0%, precision of 80.0%, and ROC-AUC of 75.8%, respectively, surpassing the best baseline models by about 5% in accuracy and 8% in precision. Other GNN models, including GAT and GCN, also demonstrated comparable performance.

Among the graph pooling layers we tested [5], the Self-Attention Graph Pooling (SAGPooling) [19] stands out to be the most effective pooling layer. Notably, the combination of GAT with SAGPooling, utilizing the AAC+QSOrder feature combination, exhibits the best performance. When paired with the GraphSAGE model, it achieved the highest metrics, with an accuracy of 79.6%, recall of 78.0%, F-1 score of 79.0%, precision of 80.0%, and ROC-AUC of 75.8%. These results significantly surpassed those of previous methods.

Other pooling layers, such as EdgePooling and ASAPooling, also showed significant improvements with the GAT and GraphSAGE models. However, GCNs did not demonstrate significant enhancement when combined with any of the pooling layers.

#### 4.2.2 The effect of Graph Size on the model performance

To test the effect of various neighborhood sizes on the model performance, we assessed the model’s performance across neighborhood graph sizes ranging from 10 to 100. The accuracy trends are depicted in Figure 3.

**Figure 3:**
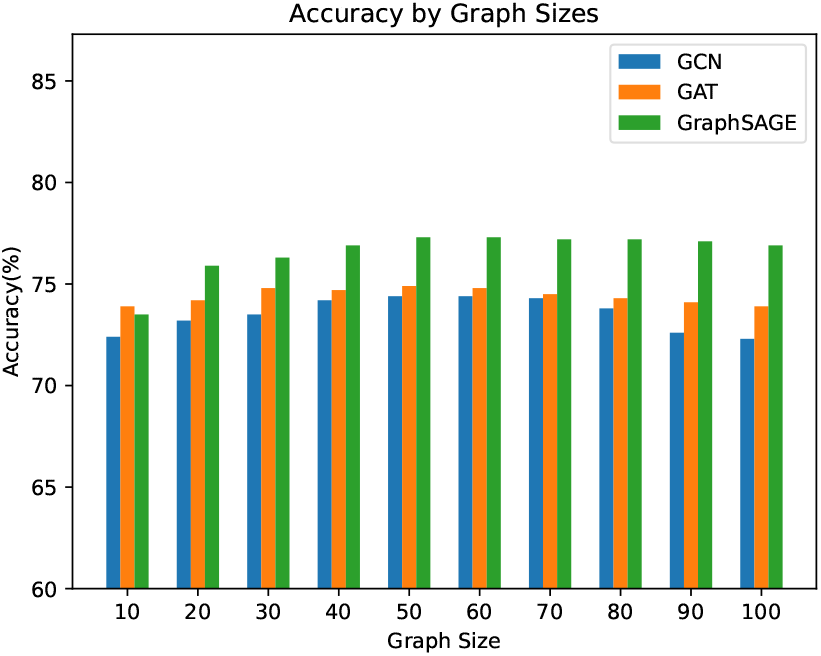
**The accuracy of the Local GNN models on neighborhood graphs of sizes 10-100. We also observed similar trends for other metrics, including F-1, recall, precision, and AUC-ROC. We can see that the model performance is approximately maximized when the graph size is between 50-60. This trend is consistent across different GNN models**

We can see that the model performance peaks at a size of 50, and marginally declines when the graph size is set to 30. For smaller graph sizes, performance significantly decreases by about 2% to 3%. Conversely, increasing the graph size to 60, and then further to 100, results in a gradual reduction in model performance. Thus, empirically, neighborhood size of 50 appears a good practical choice.

This observation likely underscores the trade-off between graph noise and information content. With small graph sizes, there is insufficient encapsulation of local patterns and interactions, leading to underperformance. Conversely, larger neihborhood sizes introduce a higher level of noise, detracting from model effectiveness.

## 5 DISCUSSION

We see that GNN models successfully combine the PPI data and physicochemical features of proteins and could distinguish MPs from non-MPs.

Across the experiments, global MP-GNN exhibited stronger performance than the local MP-GNN method. We have also run the local MP-GNN models using only on the data of *homo sapiens*, but found that the performance did not match that of the global MP-GNN.

For complexity, each single local MP-GNN with a size of 50 occupied space less than 500KB, and each training and evaluation took between 0.5-2 hours to run on a standard 3070 machine with 1 CUDA available. The full PPI network of homo sapiens was about 601MB, and the whole training and evaluation took between 15-20 hours to complete.

The accuracy difference between the global and local MP-GNNs suggests that the position of MPs in the broader graph is significant. In further work, the behaviors of MPs and non-MPs in the global and local PPI networks can be further explored. We evaluated the clustering coefficients of the local PPI graphs of the 174 MPs and 136 non-MPs. As an example, we applied statistical analysis to show that MPs (0.38 ± 0.21) exhibited higher mean clustering coefficients than non-MPs (0.27 ± 0.24), with *p <* 0.01. This suggests that the MPs are more likely to form cliques with their neighbors in the PPI networks, while non-MPs are less likely to do so.

That the GraphSAGE model outperformed the GAT model is possibly an influence of noise and small size of labelled samples for the problem. In such noisy scenarios, with possibly incorrect labels, GAT is known to perform poorly [24]. The performance did not change appreciably when we tried types of attention mechanisms – including cardinality preservation [34] and regularization [24]. This circumstance may change as more moonlighting proteins labels (and non-moonlighting proteins) become known (possibly with help of this and similar works).

We investigated the edge weights in GAT models of local PPI networks. We found that the GAT model puts large weights on the self loops – presented as larger values along the diagonal. A couple of examples of this phenomenon is shown in Figure 4. This pattern suggests that the model puts an emphasis on the presevation of individual features of nodes in the neighborhood. Thus, it would appear that this distribution of physiochemical features within the neighborhoods is important to the GAT neural network.

**Figure 4:**
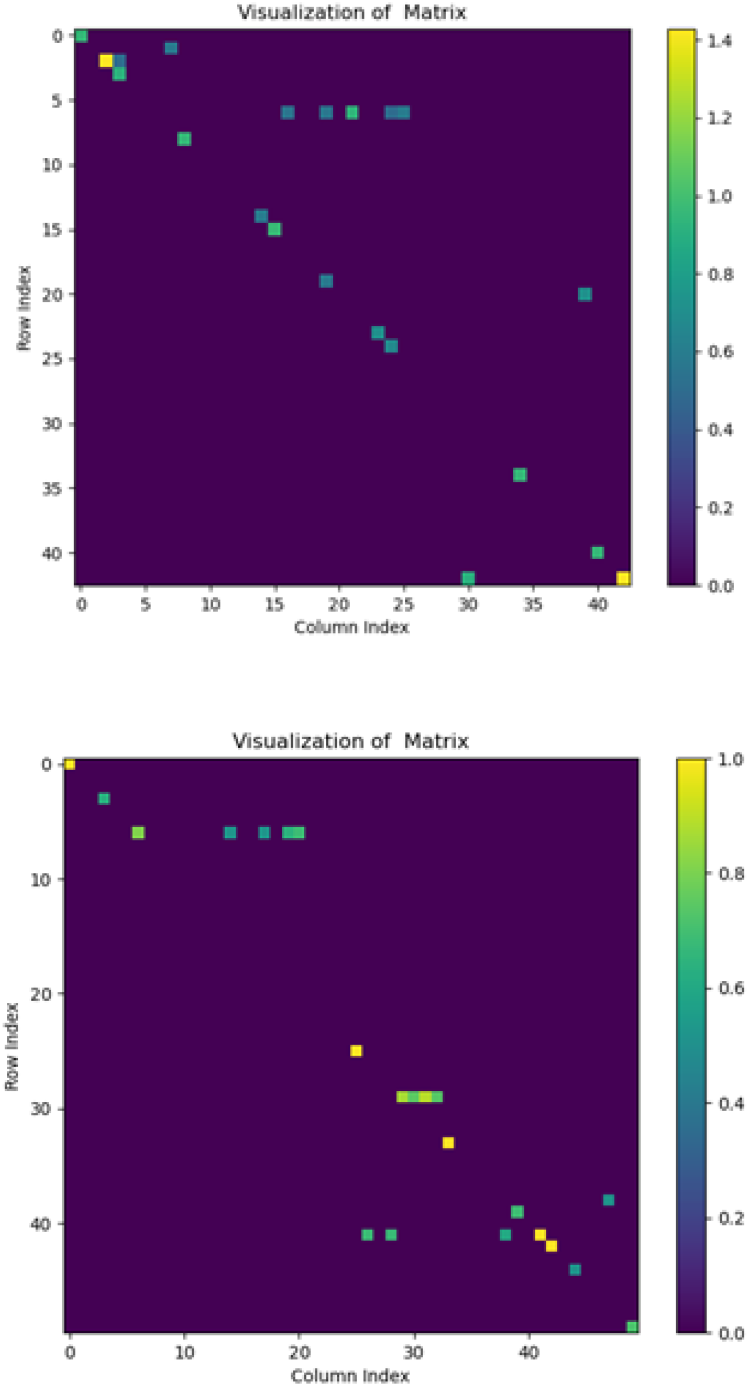
**A comparison between the Graph Attention Layer weights of MPs(top) and nonMPs(bottom) of the attention weights of two random samples. The figures were filtered with a threshold of 0.4, highlighting only those with importance. We can see that most of the important edges are centred along the diagonal line.**

## 6 CONCLUSION

There are several fronts on which this line of research can make progress. We have noted above that the data available on protein properties is relatively sparse. This is a somewhat chicken-and-egg problem. We hope that methods such as these will aid in identifying new moonlighting proteins and more labels will allow for better, more accurate models.

On the computational and statistical side, *Positive unlabelled learning* – a successor of semi-supervised learning used here – is an interesting topic in current development that can push the state of the art in this area. As can newer generations of GNNs and feature engineering.

## Footnotes

1 Note that it is hard to guarantee that any particular protein is defnitely nonmoonlighting, however, these examples are widely believed to be singe-function.

## Notes

https://github.com/VarianZhou/MPGNN.git

### Competing Interest Statement

The authors have declared no competing interest.

### Summary of Updates

Reformatted for submission to ACM-BCB 2024; Included the experiments on global PPI networks; Distinguished between local and global MP-GNNs

